# CoREST has a conserved role in facilitating SPR-5/LSD1 maternal reprogramming of histone methylation

**DOI:** 10.1101/2021.05.17.444472

**Authors:** Brandon S. Carpenter, Alyssa Scott, Robert Goldin, Sindy R. Chavez, Dexter A. Myrick, Marcus Curlee, Karen Schmeichel, David J. Katz

## Abstract

Maternal reprogramming of histone methylation is critical for reestablishing totipotency in the zygote, but how histone modifying enzymes are regulated during maternal reprogramming is not well characterized. To address this gap, we asked whether maternal reprogramming by the H3K4me1/2 demethylase SPR-5/LSD1/KDM1A, is regulated by the co-repressor protein, SPR-1/CoREST in *C. elegans* and mice. In *C. elegans*, SPR-5 functions as part of a reprogramming switch together with the H3K9 methyltransferase MET-2. By examining germline development, fertility and gene expression in double mutants between *spr-1* and *met-2*, we find that *spr-1* mutants are partially compromised for *spr-5; met-2* reprogramming. In mice, we generated a separation of function *Lsd1* M448V point mutation that compromises CoREST binding, but only slightly affects LSD1 demethylase activity. When maternal LSD1 in the oocyte is derived exclusively from this allele, the progeny phenocopy the increased perinatal lethality that we previously observed when LSD1 was reduced maternally. Together, these data are consistent with CoREST having a conserved function in facilitating maternal LSD1 epigenetic reprogramming.

## INTRODUCTION

Re-establishing the transcriptional ground state to enable embryonic development in a newly formed zygote requires extensive maternal reprogramming of chromatin at fertilization (Burton and Torres-Padilla, 2014; Hemberger et al., 2009; Li, 2002; Morgan et al., 2005; Seisenberger et al., 2012). Maternal reprogramming of chromatin is accomplished by the deposition of enzymes into the oocyte that covalently modify histones. The combination of these histone modifications contributes to developmental cell fates by regulating the accessibility of chromatin for transcription. For example, methylation of either lysine 9 or 27 on histone H3 (H3K9me and H3K27me) is generally associated with repressed transcription, whereas methylation of either lysine 4 or 36 on histone H3 (H3K4me and H3K36me) is associated with active transcription (Bannister et al., 2005; Barski et al., 2007; Bernstein et al., 2002; Bernstein et al., 2005).

Accumulating evidence suggests that patterns of histone modifications can be maintained through cell divisions to help maintain cell fate. For example, during early patterning in *Drosophila*, the expression of homeotic genes is modulated by segmentation transcription factors. After the segmentation factors turn over, the continued maintenance of homeotic gene expression through development is dependent on the H3K27 and H3K4 methyltransferases, Polycomb and Trithorax (Coleman and Struhl, 2017; Moehrle and Paro, 1994; Simon and Tamkun, 2002). Likewise, in *C. elegans*, the Polycomb Repressive Complex 2 (PRC2), which includes MES-2/3/6, maintains paternally inherited H3K27me3 during embryogenesis (Gaydos et al., 2014; Kaneshiro et al., 2019; Tabuchi et al., 2018). In addition to H3K27me3, the maternally deposited H3K36 methyltransferase, MES-4, maintains H3K36me2/3 at a subset of germline genes (MES-4 germline genes) in a transcriptionally independent manner to help reestablish the germline in the next generation (Furuhashi et al., 2010; Rechtsteiner et al., 2010). Furthermore, the transgenerational inheritance of repressive histone modifications occurs in *C. elegans* mutants lacking the COMPASS complex component, WDR-5. *wdr-5* mutants transgenerationally extend lifespan due to the accumulation of H3K9me2 across generations (Greer et al., 2010; Lee et al., 2019). Histone methylation may also be transmitted across generations in vertebrates (Brykczynska et al., 2010; Hammoud et al., 2009; Wu et al., 2011; Zheng et al., 2016; Zhu et al., 2019). Together these findings suggest that inherited histone methylation patterns are conserved across multiple phyla, and that the inheritance of the proper chromatin state is critical for normal function of the offspring.

Despite the importance of inherited histone methylation in contributing to the maintenance of cell fates, there are developmental transitions where the inheritance of histone methylation may need to be prevented. For example, in *C. elegans*, the histone demethylase SPR-5, must remove H3K4me1/2 at fertilization to prevent the inappropriate inheritance of previously specified transcriptional states. Failure to erase H3K4me1/2 at fertilization between generations in *spr-5* mutants correlates with an accumulation of H3K4me2 and ectopic spermatogenesis gene expression across ~30 generations. This accumulation of H3K4me2 leads to progressively increasing sterility, which is defined as germline mortality (Katz et al., 2009). SPR-5 reprogramming at fertilization functions together with the addition of H3K9 methylation by the methyltransferase MET-2 (Greer et al., 2014; Kerr et al., 2014). *spr-5; met-2* mutants have a synergistic sterility where progeny are sterile in a single generation, rather than over many generations (Kerr et al., 2014). Together this work supports a model in which SPR-5 and MET-2 are maternally deposited into the oocyte, where they reprogram histone methylation at fertilization to prevent defects caused by inappropriately inherited transcriptional states. Furthermore, in *C. elegans* the transgenerational maintenance of H3K36me3 by MES-4 functions to antagonize SPR-5/MET-2 repression, enabling the proper specification of the germline in the progeny (Carpenter et al., 2021).

SPR-5/MET-2 epigenetic reprogramming at fertilization is conserved in mammals. When the *met-2* ortholog *Setdb1* is maternally deleted in mice, zygotes develop slowly and die by the blastocyst stage (Eymery et al., 2016; Kim et al., 2016). Similarly, when the *spr-5* ortholog *Lsd1* is maternally deleted in mice, embryos die at the 1-2 cell stage, and these mutants are more transcriptionally similar to an oocyte than a wild type 1-2 cell embryo (Ancelin et al., 2016; Wasson et al., 2016). Thus, without maternally provided LSD1, embryos are unable to undergo the maternal-to-zygotic transition. Moreover, when maternal LSD1 protein levels are decreased, some animals can bypass the 1-2 cell arrest and survive until birth. However, progeny that are born exhibit lethality shortly after birth (perinatal lethal), indicating that incomplete reprogramming at fertilization can have phenotypes that manifest postnatally (Wasson et al., 2016). The 1-2 cell arrest and perinatal lethality phenotypes potentially occur through the inappropriate inheritance of histone methylation.

Although the evidence for maternal reprogramming across multiple taxa is mounting, it is not clear how histone modifying enzymes like LSD1 are regulated during this process. Recently, studies have implicated the co-repressor CoREST in broadly regulating LSD1 function. In mice, LSD1 and CoREST are often found in the same transcriptional corepressor complex together (Hakimi et al., 2003; Humphrey et al., 2001; Shi et al., 2003; You et al., 2001). In addition, LSD1 and CoREST have been co-crystallized, (Yang et al., 2006) and CoREST is required for the stability of LSD1 (Foster et al., 2010; Shi et al., 2005). CoREST may also be required for full LSD1 function, because although LSD1 can demethylate H3K4 peptides or bulk histones *in vitro*, it is only capable of demethylating nucleosomes when in complex with CoREST (Lee et al., 2005; Shi et al., 2005; Yang et al., 2006). Additionally, LSD1 and CoREST phenocopy each other in multiple organisms. In *Drosophila*, LSD1 and CoREST have overlapping functions in spermatogenesis and in ovary follicle progenitor cells (Lee and Spradling, 2014; Mačinković et al., 2019). In *C. elegans*, the homolog of CoREST, SPR-1, was identified, along with SPR-5, in a suppressor screen for the ability to rescue the egg-laying defect (Egl) associated with loss of SEL-12, a presenilin protein (Wen et al., 2000). Furthermore, SPR-1 has been shown to physically interact with SPR-*5 in vitro* and *in vivo* (Eimer et al., 2002; Kim et al., 2018). Together, these data raise the possibility that LSD1 could be functioning through CoREST to maternally reprogram histone methylation, but this hypothesis has not yet been tested.

Here, we utilize both mouse and *C. elegans* to test whether LSD1 and CoREST function together maternally. We demonstrate that *C. elegans* lacking SPR-1 display a reduction in brood size that is between wildtype and *spr-5* mutants. Unlike *spr-5* mutants, *spr-1* mutants do not become increasingly sterile across ~30 generations. However, when maternal reprogramming is sensitized by loss of the *met-2* gene, *met-2; spr-1* mutants reveal intermediate sterility and gene expression changes that are exacerbated compared to single mutants, but less affected than *spr-5; met-2* mutants. We also demonstrate that *met-2; spr-1* mutants misexpress MES-4 germline genes in somatic tissues at intermediate levels compared to *spr-5; met-2* mutants. In mice, we find that LSD1 and CoREST are both expressed in mouse oocyte nuclei. In addition, we generated a separation of function *Lsd1* point mutation that compromises CoREST binding, but only slightly affects LSD1 demethylase activity. When this mutation is inherited maternally, the progeny phenocopy the increased perinatal lethality that we previously observed when LSD1 was reduced maternally. Together, these data are consistent with CoREST having a conserved function in facilitating maternal LSD1 epigenetic reprogramming.

## RESULTS

### *spr-1* mutants have reduced fertility but do not exhibit germline mortality

Previously, we demonstrated that populations of *spr-5* mutants become increasingly sterile over ~30 generations (Katz et al., 2009). Therefore, if SPR-1 is required for SPR-5 maternal reprogramming activity, it is possible that *spr-1* mutants might phenocopy the germline mortality across generations observed in *spr-5* mutants. To address this possibility, we performed germline mortality assays on wild type (Bristol N2 strain, hereafter referred to as wild type), *spr-1* mutant, and *spr-5* mutant animals (Fig. 1A). Wild type hermaphrodites gave rise to ~300 progeny in the first generation (F1) and this average number of progeny was maintained through 50 generations (Fig. 1A). Similar to what we previously reported, progeny from F1 *spr-5* mutants average ~150-200 progeny, and by generation 23 (F23), the average number of progeny declined to ~60 animals (Fig. 1A). *spr-1* mutants averaged ~250 progeny in the first generation. This average number of progeny is intermediate between *spr-5* mutants and wild type. But unlike *spr-5* mutants, *spr-1* mutants never became sterile across generations (Fig. 1A).

**Figure 1.**
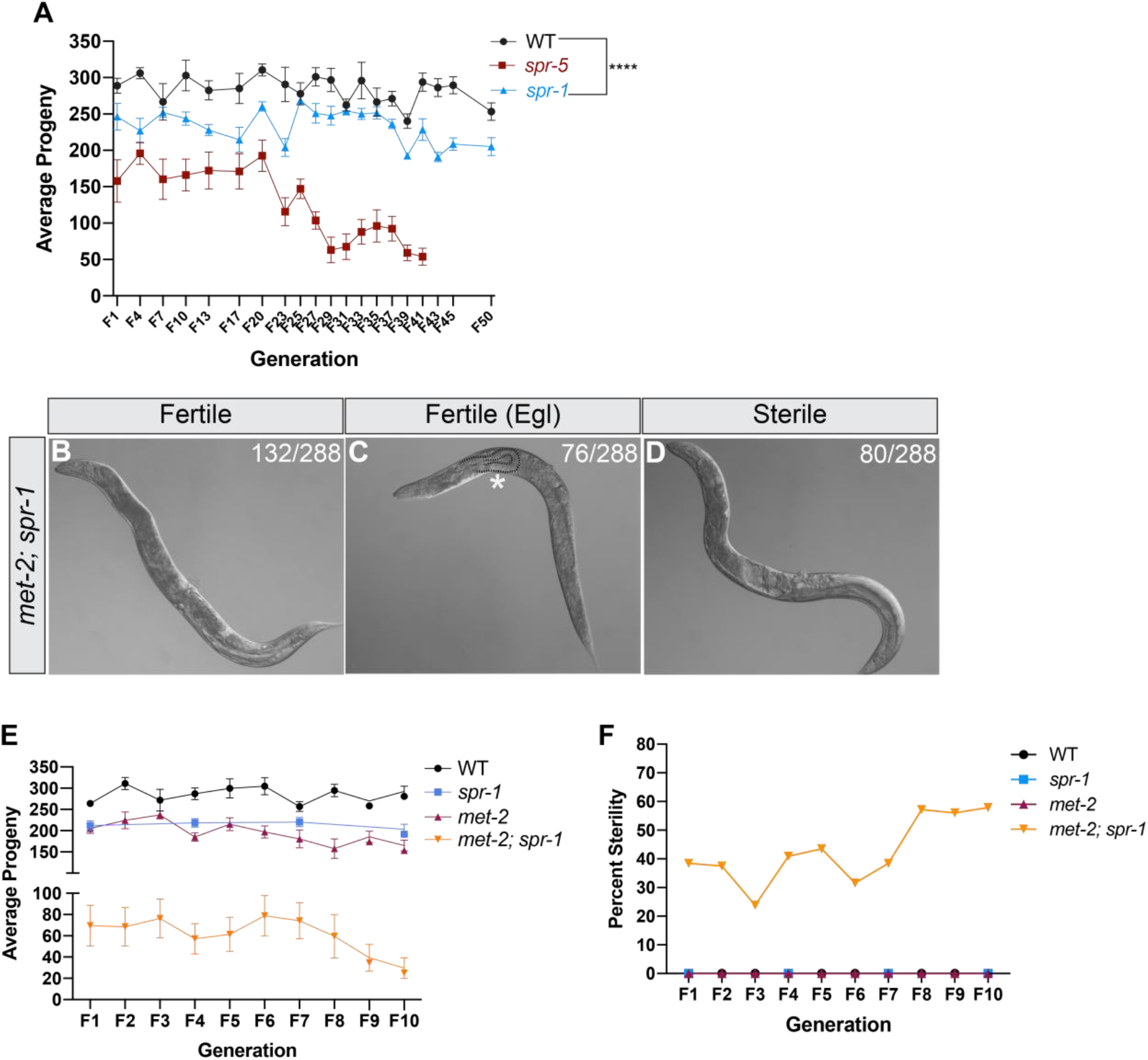
Germline mortality in *spr-1* and *met-2; spr-1* mutants. (A) The average number of total progeny from wild type (WT), *spr-5*, and *spr-1* mutants over progressive generations. The average number of progeny from *spr-1* mutants (N=192, N= total number broods counted) was significantly decreased compared to WT animals (N=92) across 50 generations (unpaired student t-test, **** p-value <0.0001). 10x DIC images of first generation (F1) *met-2; spr-1* mutants scored as either fertile (B), fertile (Egl) (C), or sterile (D). Asterisk denotes hatched larvae outlined by a dashed line inside of a *met-2; spr-1* mutant scored as fertile (Egl)(C). (E) The average number of total progeny from WT, *spr-1, met-2* and *met-2; spr-1* mutants over progressive generations. Error bars in (A, E) represent the standard error of the mean (SEM). (F) Percent of animals cloned out for experiment in (E) scored for sterility over progressive generations. *spr-1* mutant progeny were only scored at F1, F4, F7, and F10 generations in (E, F).

### *met-2; spr-1* double mutants exhibit germline mortality

Previously we demonstrated that SPR-5 synergizes with the H3K9me2 methyltransferase, MET-2, to regulate maternal epigenetic reprogramming (Kerr et al., 2014). Progeny of mutants lacking both SPR-5 and MET-2 are completely sterile in a single generation (Kerr et al., 2014). If SPR-1 is partially required for SPR-5 maternal reprogramming, then it is possible that *spr-1* mutants will exhibit a synergistic phenotype when combined with a *met-2* mutation. To determine whether *spr-1* mutants display any abnormal phenotypes in a *met-2* mutant background, we individually cloned out 288 progeny from first generation (F1) *met-2; spr-1* double mutants and examined each animal for sterility (Fig. 1B-D). Of the 288 individually cloned *met-2; spr-1* F1 mutant progeny, 208 were fertile (Fig. 1B,C) and 80 were sterile (Fig. 1D). We also observed that 76 of the 208 fertile F1 progeny die as young adults due to defects in egg laying (Fig. 1C). To determine whether fertile *met-2; spr-1* mutants become germline mortal, we counted the average number of progeny from *met-2; spr-1* mutants over successive generations and compared them to wild type, *spr-1* and *met-2* mutants (Fig. 1E; Fig. S1A,C). As observed previously, the average number of progeny from *spr-1* and *met-2* mutants were lower than wild type, but remained consistent over 10 generations, and neither mutant gave rise to sterile animals over that time frame (Fig. 1E,F; Fig. S1A,B). Fertile *met-2; spr-1* mutant progeny produced an average of ~70 progeny in the first generation. However, by generation 10 the average number of progeny declined to ~30 (Fig. 1E; Fig. S1C). Consistent with this germline mortality phenotype, the number of completely sterile animals in *met-2; spr-1* mutants increased across successive generations from ~30% at early generations to ~60% by F10 (Fig. 1F; Fig. S1D). Together, these results show that *met-2; spr-1* mutants display a germline mortality phenotype that is intermediate between the maternal effect sterility of *spr-5; met-2* mutants in a single generation and the germline mortality of *spr-5* and *met-2* single mutants over ~30 generations.

### The sterility of *met-2; spr-1* mutants resembles *spr-5; met-2* mutants

We also examined the gonads of sterile *met-2; spr-1* mutants to determine if the sterility resembles the sterility of *spr-5; met-2* mutants. *spr-5; met-2* mutants have a squat germline, with both gonad arms failing to elongate (Fig. S2A,B; Kerr et al., 2014). Within the squat germline there are proliferating germ cells, sperm and oocytes, indicating that the germline has proceeded through normal transitions. However, these cell types are inappropriately interspersed (Carpenter et al., 2021; Katz et al., 2009). Unlike *spr-5; met-2* mutants, F1 sterile *met-2; spr-1* mutants have elongated gonad arms. However, within these sterile F1 gonads, we observed a similarly disorganized mixture of germ cells, sperm and oocytes (Fig. S2C,D). In addition, after several generations the germlines of sterile *met-2; spr-1* mutants resembled the squat germlines of *spr-5; met-2* mutants, although unlike *spr-5; met-2* mutants, some animals remained partially fertile at later generations (Fig. 1E; Fig. S2A,B,E,F). Thus, the sterility of *met-2; spr-1* mutants at late generations phenocopies the maternal effect sterility of *spr-5; met-2* mutants observed in the first generation.

### Transcriptional misregulation in *met-2; spr-1* progeny resembles that observed in *spr-5; met-2* progeny but is less affected

Since the severity of the germline mortality phenotype of *met-2; spr-1* mutants is between *spr-5; met-2* mutants and *spr-1* or *met-2* single mutants, it raises the possibility that maternal SPR-5 reprogramming may be partially dependent upon the SPR-5 interacting partner SPR-1. If mutating *spr-1* partially compromises SPR-5 maternal reprogramming, we would expect that the genes that are misexpressed in *met-2; spr-1* mutants would be similar to *spr-5; met-2* mutants, but that the gene expression changes would be less affected in *met-2; spr-1* mutants. To test this possibility, we performed RNA-seq on F7 *spr-1, met-2*, and *met-2; spr-1* mutant L1 progeny compared to wild type L1 progeny. We chose to perform the analysis on F7 *met-2; spr-1* mutants because this generation precedes the increase in sterility that we observed in our germline mortality assay after F7 (Fig. 1F; Fig. S1A). Thus, by performing the analysis at F7, it allowed us to observe primary effects from the loss of MET-2 and SPR-1, rather than secondary effects due to the sterility. In addition, we utilized starved L1 larvae for our RNA-seq analysis for two reasons. First, starved L1 larvae only have two germ cells that are not undergoing transcription. As a result, performing RNA-seq on these larvae allows us to exclusively examine somatic transcription. Second, we have previously performed RNA-seq and differential gene expression analysis on the L1 stage of *spr-5; met-2* mutant progeny, so performing the RNA-seq analysis on *met-2; spr-1* mutants at the L1 stage allows us to compare to our previously published data set (Carpenter et al., 2021). We identified 1,787 differentially expressed genes (DEGs) in *met-2; spr-1* mutant progeny compared to wild type (Fig. S3A; Fig. S4C,F), and most of these genes are differentially expressed in *met-2* (856/1327)(Fig. S3A,B; Fig. S4B,E) and *spr-1* single mutants (40/60)(Fig. S3A,B; Fig. S4A,D) compared to wild type (Fig. S3B).

To determine whether gene expression changes in *met-2; spr-1* mutants resemble those in our previously published *spr-5; met-2* mutant RNA-seq data set, we first compared DEGs between the two data sets. We identified 1,787 DEGs in *met-2; spr-1* mutant progeny compared to wild-type. 1,010 (57%) of these significantly overlapped with the 4,223 DEGs that we previously identified in *spr-5; met-2* mutant progeny compared to wild-type (Fig. 2A, hypergeometric test, P-value < 1.28E-270, (Carpenter et al., 2021), and the gene ontology categories of DEGs in both data sets are similar (Fig. S3C,D; Carpenter et al., 2021). We also examined the overlapped gene expression changes between up-regulated and down-regulated DEGs in both datasets separately. Of the 1,067 up-regulated DEGs in *met-2; spr-1* mutant progeny, 676 (63%) of these overlap with the 2,330 up-regulated DEGs in *spr-5*; *met-2* mutant progeny (Fig. 2B, hypergeometric test, P-value < 2.61E-392; Carpenter et al., 2021). Of the 720 down-regulated DEGs in *met-2; spr-1* mutant progeny, 236 (33%) of these overlap with the 1,893 DEGs down-regulated DEGs in *spr-5*; *met-2* mutant progeny (Fig. 2C, hypergeometric test, P-value < 2.16E-72; Carpenter et al., 2021).

**Figure 2.**
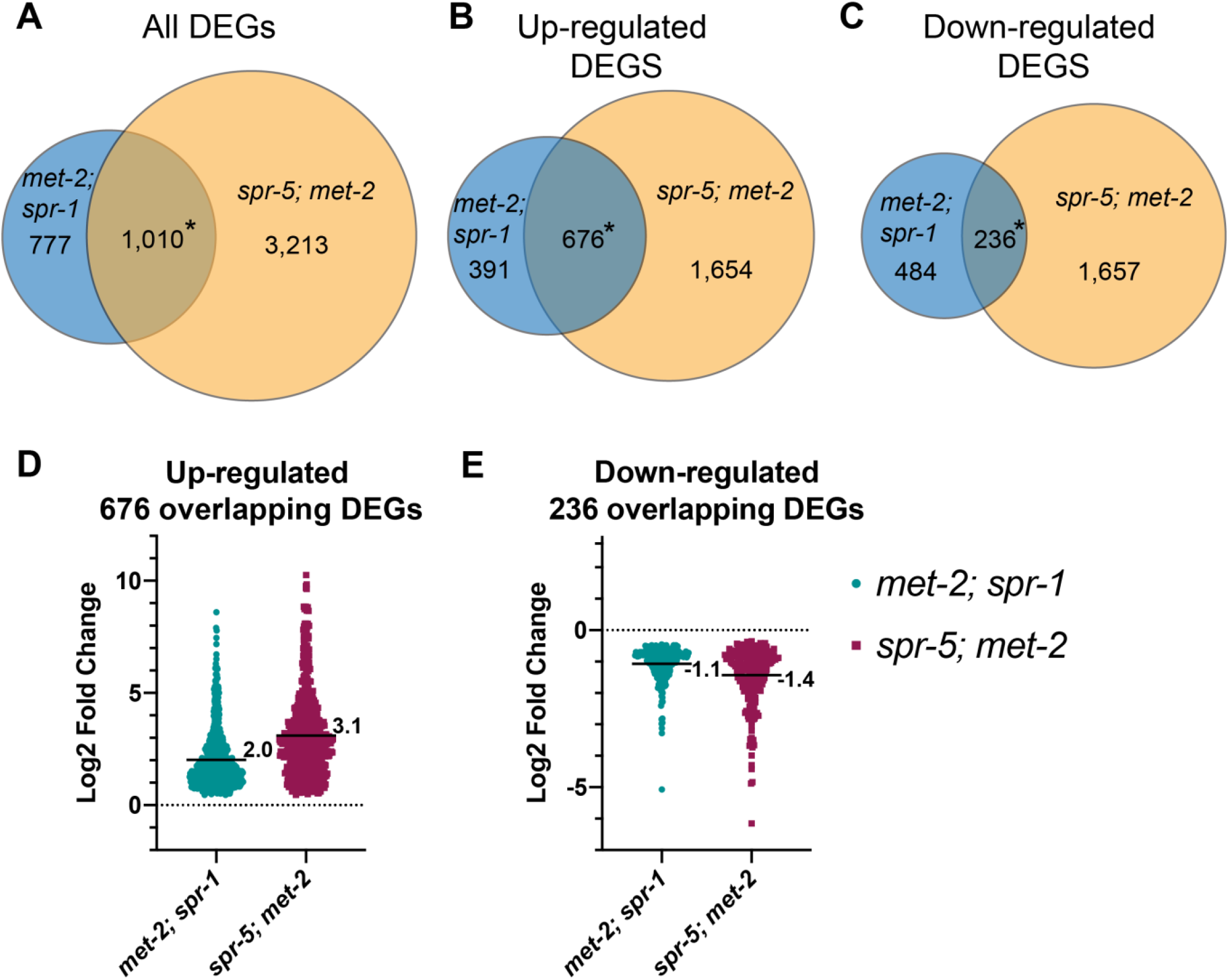
Transcriptional misregulation in *met-2; spr-1* progeny resembles that observed in *spr-5; met-2* progeny but is less affected. Overlap between all (A), up-regulated (B), and down-regulated (C) differentially expressed genes (DEGs) in *met-2; spr-1* and *spr-5; met-2* L1 progeny. Significant over-enrichment in A-C was determined by the hypergeometric test (*P-value < 1.28E-270, *P-value < 2.61E-392, *P-value < 2.16E-72, respectively). Scatter dot plots displaying the log2 fold change of the 676 up-regulated (D), and 236 down-regulated (E) overlapping DEGs between *met-2; spr-1* and *spr-5; met-2* progeny. (D, E) Numbers and solid black lines represent the mean log2 fold change. DEGs in *spr-5; met-2* progeny were obtained from (Carpenter et al., 2021).

If mutating *spr-1* partially compromises SPR-5 maternal reprogramming, we would expect that the gene expression changes in *met-2; spr-1* mutants would be less affected in *spr-5*; *met-2* mutants. To determine if this is the case, we compared the average log2 fold change (FC) of the upregulated and downregulated DEGs separately. The average log2 (FC) of DEGs that are up-regulated in *met-2; spr-1* mutant progeny is 2, compared to 3.1 in *spr-5; met-2* mutant progeny (Fig. 3D; Carpenter et al., 2021). Similarly, the average log2(FC) of DEGs that are down-regulated in *met-2; spr-1* mutant progeny is −1, compared to −1.4 in *spr-5; met-2* mutant progeny (Fig. 3E; Carpenter et al., 2021). Together, these data demonstrate while the same genes are differentially expressed in *spr-5; met-2* and *met-2; spr-1* mutant progeny, the changes are smaller in *met-2; spr-1* mutant progeny.

**Figure 3.**
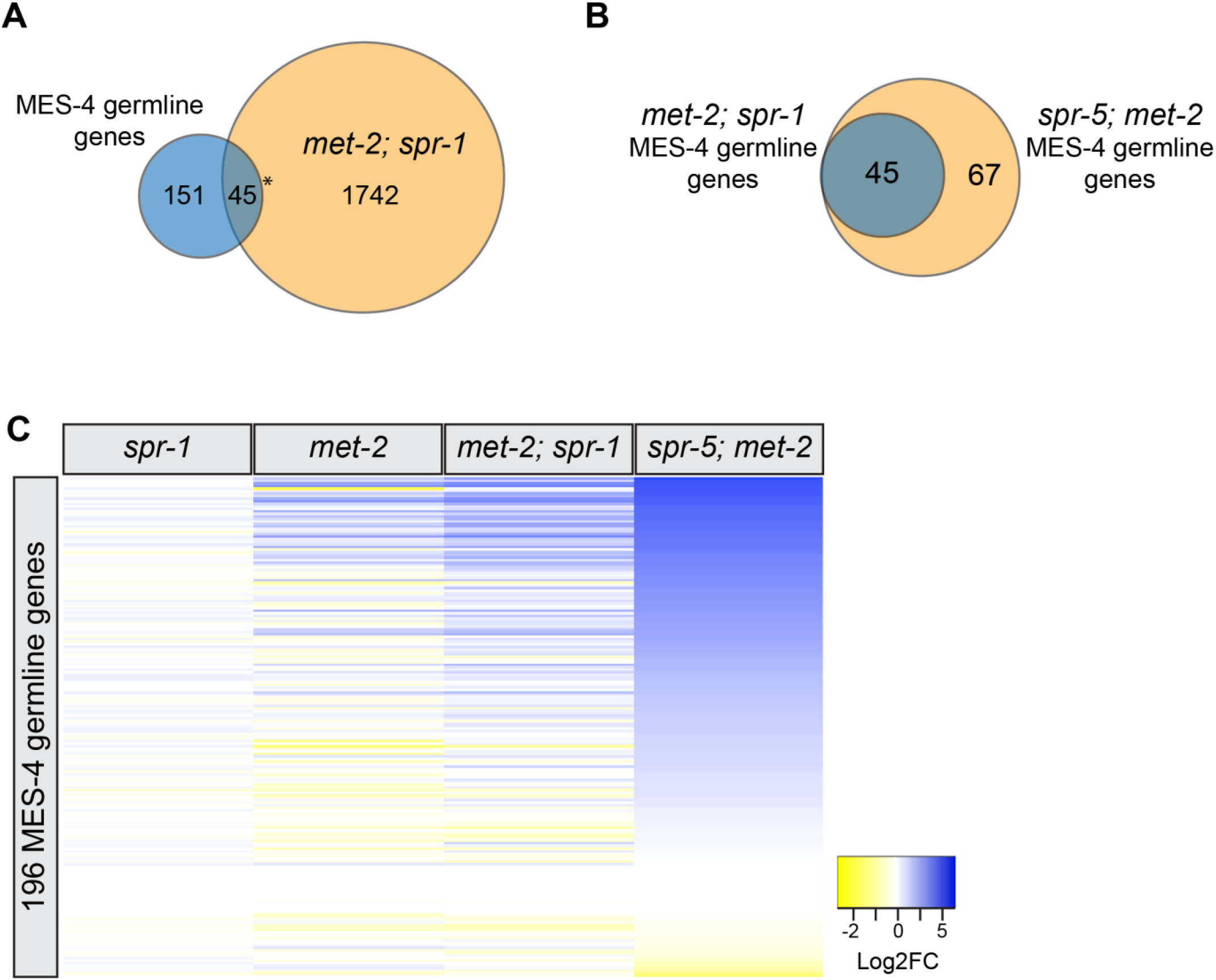
MES-4 germline genes are enriched in *met-2; spr-1* mutants, but less affected compared to *spr-5; met-2* mutants. (A) Overlap between MES-4 germline genes and differentially expressed genes (DEGs) in *met-2; spr-1* L1 progeny. Asterisks denotes significant over-enrichment in A as determined by a hypergeometric test (P-value < 1.41E-9). (B) Overlap between MES-4 germline genes differentially expressed in *met-2; spr-1* and *spr-5; met-2* L1 progeny. (C) Heatmap of log2 fold change (FC) of all 196 MES-4 germline genes in *spr-1, met-2, met-2; spr-1* and *spr-5; met-2* mutants compared to wild type. log2(FC) values are represented in a yellow to blue gradient with a range of −2 to 5. Yellow represents genes with negative log2(FC) values and blue represents genes with positive log2FC values compared to wild type.

### MES-4 germline genes are enriched in *met-2; spr-1* mutants, but less affected compared to *spr-5; met-2* mutants

MES-4 germline genes are genes that are expressed in the parental germline and acquire H3K36 methylation, which is maintained by a transcription independent methyltransferase MES-4 in the embryo of the progeny. Recently, we demonstrated that *spr-5; met-2* mutants ectopically express 112 (57%) out of the 196 MES-4 germline genes in somatic tissues (Carpenter et al., 2021). If the loss of SPR-1 partially compromises SPR-5 function, we would expect that MES-4 germline genes would also be ectopically expressed in *met-2; spr-1* mutants, though to a lesser degree. Of 196 MES-4 germline genes, 45 (23%) MES-4 germline genes were misexpressed in *met-2; spr-1* mutant progeny compared to wild type (Fig. 3A, hypergeometric test, P-value < 1.41E-9). All of these 45 MES-4 germline genes overlap with the 112 MES-4 germline genes that are misregulated in *spr-5; met-2* mutants (Fig. 3B; Carpenter et al., 2021). Thus, like *spr-5; met-2* mutants, *met-2; spr-1* mutants ectopically express MES-4 germline genes. However, when we compared the log2(FC) in expression of all of the MES-4 germline genes in *spr-1, met-2, met-2*; *spr-1*, and *spr-5; met-2* mutant progeny compared to wild type, we observed that the changes in the levels of gene expression are less affected in *met-2*; *spr-1* mutants than *spr-5; met-2* mutants (Fig. 3C; Carpenter et al., 2021).

### LSD1 and CoREST are expressed during each stage of mouse oocyte development

Taken together, our results are consistent with SPR-5/LSD1 functioning maternally through SPR-1/CoREST in *C. elegans*. To determine whether there is a role for CoREST in LSD1 maternal reprogramming in mammals, we also sought to investigate the maternal interaction between LSD1 and CoREST in mice. Previous studies have shown that LSD1 is expressed during all stages of mouse oocyte development (Wasson et al. 2016, Ancelin et al. 2016, Kim et al. 2015). If LSD1 and CoREST function together in a complex, we would expect them to be expressed during the same stages of oogenesis. Previously CoREST was shown to be expressed in mouse oocytes, but the precise stages of oogenesis in which CoREST is expressed were not characterized (Ma et al., 2012). Thus, to determine whether CoREST is expressed at the same time as LSD1 in mouse oogenesis, we performed immunofluorescence experiments and examined CoREST and LSD1 protein at the primary, secondary, and antral stages of oocyte development (Fig. 4). Identical to what we and others previously observed with LSD1 (Fig. 4 A-I; Ancelin et al., 2016; Kim et al., 2015b; Wasson et al., 2016), CoREST was also highly expressed in the oocyte nucleus and in the surrounding follicle cells during all stages of oocyte development (Fig. 4J-R).

**Figure 4.**
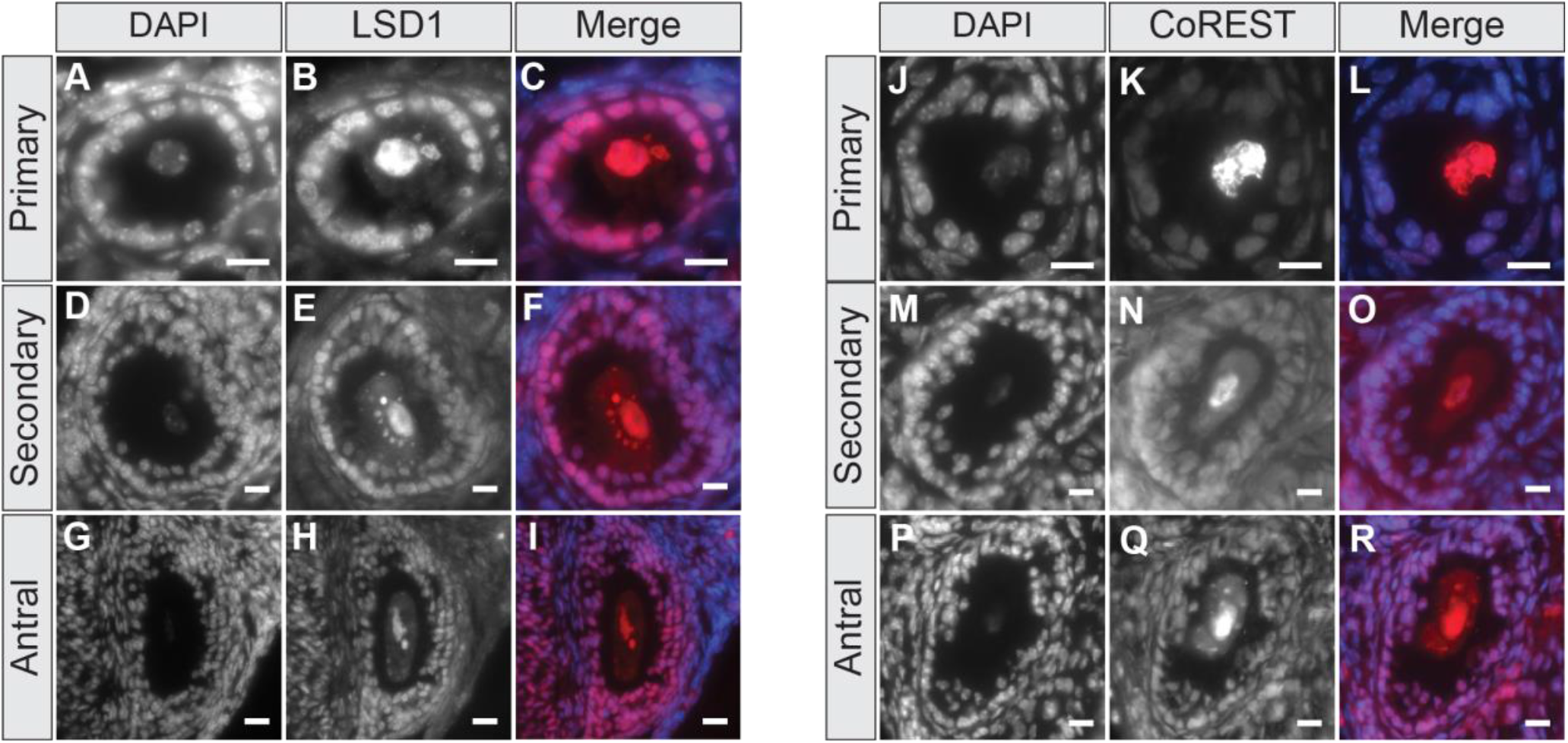
LSD1 and CoREST are expressed during each stage of mouse oocyte development. Representative immunofluorescence images of various stages of the mouse oocyte: primary (A-C, J-L), secondary (D-F, M-O), and antral (G-I, P-R). DAPI (A, D, G, J, M, P), LSD1 (B, E, H) CoREST (K, N, Q), and Merge (C, F, I, L, O, R). Both LSD1 and CoREST are expressed in the oocyte nucleus and surrounding follicle cells during each stage of oocyte development. Scale bars= 25um.

### Reducing the function of maternally-provided LSD1 causes perinatal lethality

Since CoREST has the same expression pattern as LSD1 in mouse oocytes, we wanted to test whether LSD1 functions through CoREST by specifically disrupting the presumptive CoREST-LSD1 interaction in the mouse oocyte. To do this, we utilized CRISPR to generate a point mutation, M448V, in the tower domain of the *Lsd1* gene at the endogenous locus (Fig. 5A; Fig. S5). This allele will be referred to as *Lsd1*^*M448V*^. The tower domain is the site of protein-protein interactions (Forneris et al., 2007; Stavropoulos et al., 2006; Yang et al., 2006), and M448V resides in a residue that binds CoREST (Shi et al., 2005). Previous studies have shown that this mutation reduces the ability of LSD1 to demethylate histones *in vitro* (85% demethylase activity compared to wild-type LSD1) (Nicholson et al., 2013). This modest reduction in LSD1 function is unlikely to compromise maternal reprogramming. However, the M448V mutation severely reduces the ability of LSD1 to bind CoREST (35% binding activity compared to wild type *in vitro*) (Nicholson et al., 2013). Thus, the M448V mutation serves as a separation-of-function allele between demethylase activity and CoREST binding.

**Figure 5.**
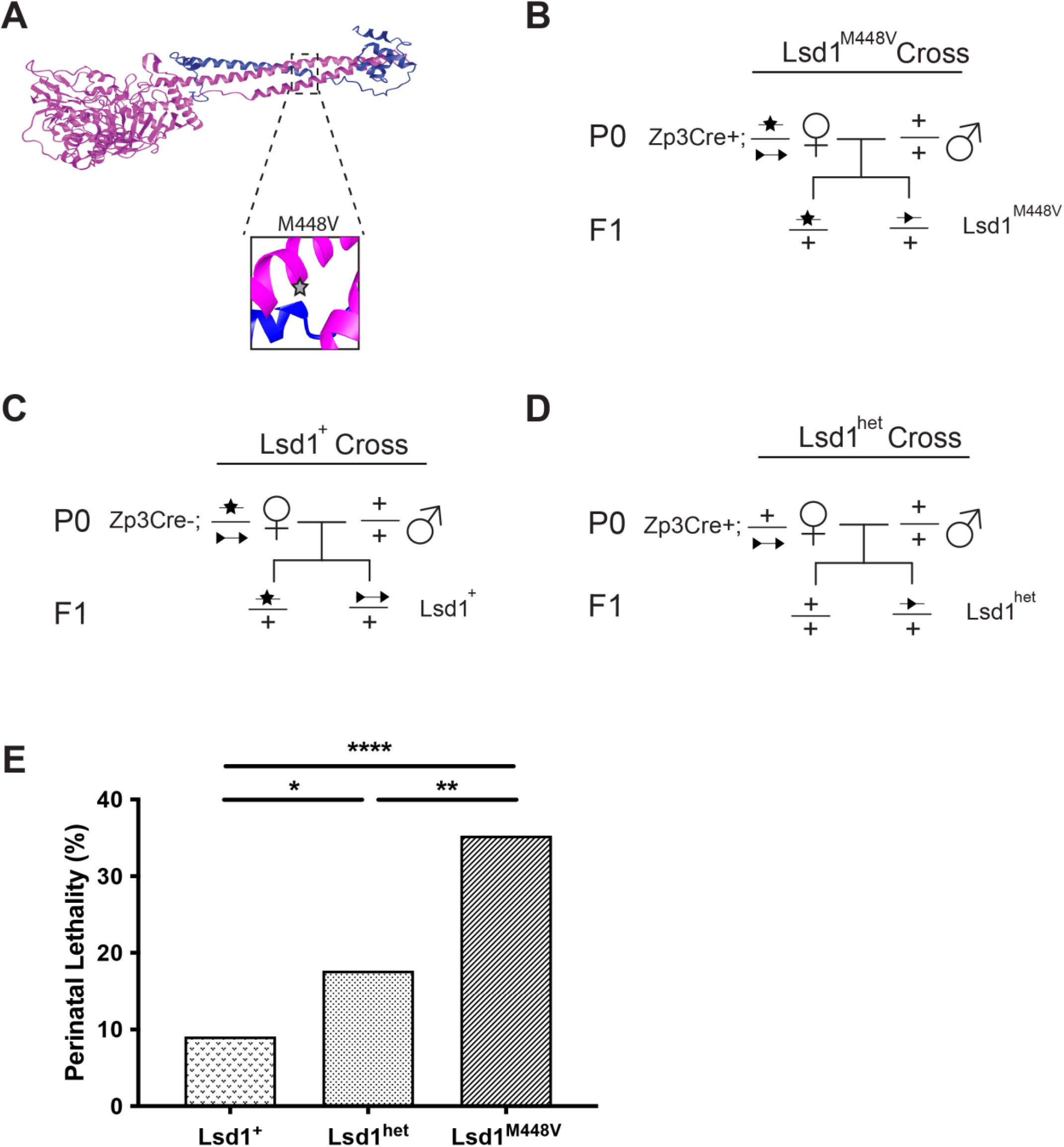
Hypomorphic maternal LSD1 results in perinatal lethality. (A) Crystal structure of LSD1 (pink) in complex with CoREST (blue) from Nicholson et al., 2013. The M448V mutation is in a CoREST binding site (star). (B-D) Genetic crosses showing wild type (+), loxP sites (triangles), and M448V (star) alleles. In all cases, P0 females are crossed to wild-type males, so that F1 progeny have normal zygotic LSD1 activity from their paternal allele after transcription begins at the 2-cell stage. (B) In the *Lsd1*^*M448V*^ cross, P0 mothers are Zp3Cre+, contributing only the hypomorphic allele maternally. (C) In the Lsd1^+^ control cross, P0 mothers are Zp3Cre-, contributing a wild-type and hypomorphic allele maternally. (D) In the Lsd1^het^ control cross, P0 mothers are Zp3Cre+, contributing one wild-type copy of *Lsd1* maternally. (E) Percent perinatal lethality per litter by experimental condition, n= 154 pups from 24 litters (Lsd1^+^), n= 96 pups from 15 litters (Lsd1^het^), and n= 187 pups from 32 litters (Lsd1^M448V^). See Supplemental file 1 for the list of individual litters. p values calculated using a chi-square test, **** = p<.0001, ** = p<.01, *= p<.05.

To interrogate the interaction between LSD1 and CoREST specifically in oocytes, we utilized our newly generated M448V *Lsd1* mutation. Previous studies have shown that a complete loss of LSD1 protein in the mouse oocyte results in embryonic arrest of offspring at the 1-2 cell stage (Ancelin et al., 2016; Wasson et al., 2016). This arrest is due to a failure to undergo the maternal-to-zygotic transition in gene expression. When LSD1 protein levels are decreased in the mouse oocyte, embryos can bypass the embryonic arrest, but ~30% of animals die perinatally, shortly after birth (Wasson et al., 2016). Our results in *C. elegans* suggest that loss of CoREST results in a partial loss of LSD1 function. Therefore, if LSD1 function partially requires CoREST maternally in mice, we would expect that having only the *Lsd1*^*M448V*^ allele maternally would result in offspring that phenocopy the perinatal lethality observed from a partial loss of maternal LSD1. To test this possibility, we generated mice with the *Lsd1*^*M448V*^ allele over a floxed allele of the *Lsd1* gene. In the presence of an oocyte-specific *Zp3-Cre* allele that expresses prior to the first meiotic division, the floxed allele recombines to a null allele in the oocyte. As a result, the only maternal contribution of LSD1 is from the *Lsd1*^*M448V*^ allele, which produces LSD1 with a reduced ability to bind CoREST (Fig. 5B). The F1 offspring from this cross will be referred to as *Lsd1*^*M448V*^ progeny. Importantly, the mothers have a normal copy of the *Lsd1* gene in every other cell type throughout the mouse, and heterozygous *Lsd1* animals have been shown to have no phenotypic defects (Engstrom et al., 2020; Foster et al., 2010; Jin et al., 2013; Wang et al., 2007). In addition, mothers with a compromised maternal *Lsd1* allele are crossed to wild-type males, so the progeny have normal zygotic LSD1 activity from their paternal allele after transcription begins at the 2-cell stage. This mating scheme enables us to determine the specific effect of compromising LSD1 activity maternally in the oocyte. In one set of controls, the mother has the *Lsd1*^*M448V*^ allele over a floxed allele of LSD1 and is *Zp3Cre* negative (Fig. 5C). The maternal contribution in this case would be one functional copy of LSD1 and one hypomorphic *Lsd1*^*M448V*^ copy. These control F1 offspring will be referred to as *Lsd1*^*+*^. In the other set of controls, the mother has a wild-type copy of the *Lsd1* gene over a floxed allele of *Lsd1*, and are *Zp3Cre* positive (Fig. 5D). The maternal contribution will be just one functional copy of the *Lsd1* gene. Previous studies have shown animals that are heterozygous for the *Lsd1* null allele have 70% protein levels (30% reduction) compared to homozygotes (Engstrom et al., 2020). These control F1 offspring will be referred to as *Lsd1*^*het*^.

If disrupting the interaction between LSD1 and CoREST specifically in the oocyte phenocopies the perinatal lethality that we previously observed when LSD1 is hypomorphic maternally, it would provide further evidence that LSD1 functions in a complex with CoREST in the oocyte. To address this possibility, we examined perinatal lethality between postnatal day 0 (P0) and P1 in *Lsd1*^*M448V*^ progeny versus *Lsd1*^*+*^ and *Lsd1*^*het*^ controls. All litters were generated from mothers that were less than 8 months old, to avoid any complications associated with advanced maternal age. Overall, we observe increasing perinatal lethality with increasingly compromised maternal LSD1. In *Lsd1*^*+*^ control progeny, when one allele of *Lsd1* lacks the ability to bind CoREST, we observe 9% (N=24) perinatal lethality during the first 48hrs after birth. When one copy of *Lsd1* is fully deleted maternally (*Lsd1*^*het*^ progeny), the perinatal lethality increased to 18% (N=15), and when maternal LSD1 is solely provided from the *Lsd1*^*M448V*^ allele, perinatal lethality further increases to 35% (N=32)(Fig. 6E). The 35% perinatal lethality, when LSD1 completely lacks the ability to bind CoREST maternally, is similar to the ~30% perinatal lethality that we previously observed when LSD1 is partially lost maternally (Wasson et al., 2016). Importantly, the level of perinatal lethality does not depend on which allele the pup inherits from its mother (Fig. S6). Moreover, the entire litter died in 10 litters of *Lsd1*^*M448V*^ progeny out of 32 total (31.2%), versus only 2 litters out of 14 (14.2%) in *Lsd1*^*het*^ animals, and 0 out of 24 litters (0%) in *Lsd1*^*+*^ controls (Supplemental file 1). The 31% of *Lsd1*^*M448V*^ litters in which all of the animals within the litter die is similar to the 10 out of 20 litters (50%) in which the entire litter died that we previously observed upon partial loss of maternal LSD1 (Wasson et al., 2016). Thus, the loss of LSD1’s ability to bind CoREST maternally phenocopies the perinatal lethality observed in progeny from mothers with partial loss of LSD1 protein in the oocyte.

## DISCUSSION

### CoREST regulates LSD1 maternal reprogramming of histone methylation

Despite our increasing knowledge of the enzymes involved in maternal epigenetic reprogramming, how these enzymes are regulated remains unclear. To begin to address this question, we asked whether maternal epigenetic reprogramming by the H3K4me1/2 demethylase SPR-5/LSD1/KDM1A is dependent on SPR-1/CoREST. In *C. elegans*, we find that the fertility of *spr-1* mutants is intermediate between *spr-5* mutants and wild type, which raises the possibility that loss of the *spr-1* gene partially compromises SPR-5 maternal reprogramming. However, *spr-1* mutants do not phenocopy the germline mortality phenotype of *spr-5* mutants. This suggests that if SPR-1 contributes to SPR-5 reprogramming, SPR-5 function is not completely dependent upon SPR-1.

Because SPR-5 and the H3K9 methyltransferase MET-2 function together in maternal reprogramming (Carpenter et al., 2021; Kerr et al., 2014; Greer et al., 2014), it provides a unique opportunity to ask whether SPR-1 functions in maternal SPR-5 reprogramming by making double mutants between *spr-1* and *met-2*. If loss of SPR-1 partially compromises SPR-5 reprogramming, then *spr-1* mutants might also display a synergistic sterility phenotype when combined with a mutation in *met-2*. Consistent with this possibility, *met-2; spr-1* double mutants had a germline mortality phenotype that is intermediate between the maternal effect sterility of *spr-5; met-2* mutants and the germline mortality of *spr-5* and *met-2* single mutants. To determine if this synergistic sterility phenotype is due to loss of SPR-1 partially compromising SPR-5 reprogramming, we performed two additional experiments. First, we examined the gonads of *met-2; spr-1* double mutants to determine if the germline phenotype resembles the germline phenotype of *spr-5; met-2* mutants. This analysis demonstrated that the germline phenotype of *met-2; spr-1* mutants at late generations, with a squat gonad and disorganized germline cell types, is similar to *spr-5; met-2* mutants. This is consistent with the possibility that loss of SPR-1 partially compromises SPR-5 reprogramming. Second, we performed RNA-seq on F7 *met-2; spr-1* mutants. If SPR-1 functions specifically with SPR-5, we would expect that the genes that are misexpressed in *met-2; spr-1* mutants to be similar to the genes that are affected in *spr-5; met-2* mutants. Consistent with this possibility we observe a significant overlap in differentially expressed genes between *met-2; spr-1* mutants and *spr-5; met-2* mutants. In addition, the gene expression pathways affected in *met-2; spr-1* mutants are similar to those affected in *spr-5; met-2* mutants. However, if loss of SPR-1 only partially compromises SPR-5 function, we would expect that the magnitude of the gene expression changes in *met-2; spr-1* mutants would be less changed than in *spr-5; met-2* mutants. Strikingly, we find that the gene expression changes in *met-2; spr-1* mutants are consistently less affected than those that we observed previously in *spr-5; met-2* mutants.

MES-4 germline genes become ectopically expressed in the absence of SPR-5/MET-2 reprogramming. If SPR-1 partially compromises maternal reprogramming, we would expect that MES-4 germline genes would also be ectopically expressed in the soma of *met-2; spr-1* mutants. The starved L1 larvae that we used for *met-2; spr-1* mutant RNA-seq only have two germ cells that are not undergoing transcription. Despite this, we observe the expression of MES-4 germline genes in L1 *met-2; spr-1* mutants. This suggests that, like we previously observed in *spr-5; met-2* mutants, *met-2; spr-1* mutants ectopically express MES-4 germline genes in somatic tissues. In addition, the fact that the MES-4 germline genes that are affected by loss of SPR-1 are the same as those affected by the loss of SPR-5 suggests that they function together on the same MES-4 targets. The ectopic expression of MES-4 germline genes in *met-2; spr-1* mutants provides further evidence that SPR-1 is functioning in maternal SPR-5 reprogramming. However, the magnitude of the ectopic expression of MES-4 genes in *met-2; spr-1* mutants is intermediate between *spr-5; met-2* double mutants and *spr-1* or *met-2* single mutants. Taken together, these data suggest that SPR-5 functions maternally through its interaction with SPR-1, with loss of SPR-1 partially compromising SPR-5 reprogramming. Consistent with this conclusion, SPR-1 and SPR-5 have been shown to directly interact with one another in *C. elegans* (Eimer et al., 2002; Kim et al., 2018), and both were identified in a screen for suppressors of presenilin (Wen et al., 2000).

In mammals, epigenetic reprogramming at fertilization also requires LSD1/SPR-5 and SETDB1/MET-2. To determine whether LSD1 reprogramming in mammals requires CoREST, we addressed the maternal role of CoREST in mice. We found that CoREST and LSD1 are both expressed during all stages of mouse oocyte development, indicating that both proteins are spatially and temporally positioned for LSD1 to be functioning through CoREST in maternal reprogramming. This is consistent with previous literature describing the expression of LSD1 (Ancelin et al., 2016; Kim et al., 2015b; Wasson et al., 2016) and CoREST (Ma et al., 2012) in the mouse oocyte. Previously, we found that decreased levels of LSD1 protein in the oocyte results in ~30% perinatal lethality in progeny derived from these mothers (Wasson et al., 2016). Here we show that a M448V mutation, that reduces the ability to bind CoREST, phenocopies the perinatal lethality phenotype observed when LSD1 maternal protein is partially decreased, including the observation that many times all of the animals in a particular litter die. This suggests that the partial requirement for CoREST in maternal LSD1 reprogramming is conserved in mammals. In addition, we detect an allelic series in which the percentage of perinatal lethality increases from *Lsd1*^*+*^ to *Lsd1*^*het*^ progeny, and increases again from *Lsd1*^*het*^ to *Lsd1*^*M448V*^ progeny. The finding that the extent of perinatal lethality is more severe in *Lsd1*^*het*^ progeny compared to *Lsd1*^*+*^ progeny, provides further evidence that the *Lsd1*^*M448V*^ allele only partially compromises maternal LSD1 activity. In addition, we find that further compromising maternal LSD1 reprogramming in *Lsd1*^*M448V*^ progeny compared to *Lsd1*^*het*^ progeny leads to a further increase in perinatal lethality. This strengthens the link that we previously observed (Wasson et al., 2016) between maternal epigenetic reprogramming and defects that manifest postnatally. However, it remains to be determined whether these defects are due to the direct inheritance of inappropriate histone methylation or due to an indirect effect through some other epigenetic mechanism.

### Evidence from diverse developmental processes across multiple phyla support a role for CoREST in LSD1 function

Across multiple phyla, the function of CoREST extends beyond the female germline. In *Drosophila*, loss of *Lsd1* results in sterility in both males and females (Reuter et al. 2007, Di Stefano et al. 2007, Szabad et al. 1988). However, CoRest and Lsd1 also function in the germline support cells. Lsd1 is required in escort cells that support early female germline differentiation (Eliazer et al., 2011), and knockdown of either Lsd1 or CoRest protein causes a number of phenotypes in ovarian follicle cells (Domanitskaya and Schüpbach, 2012; Lee and Spradling, 2014). The requirement for CoRest in cells that support oogenesis causes sterility in female CoRest mutants. Together, these data potentially implicate CoRest in regulating Lsd1 function, but the direct role of CoRest has yet to be determined during oogenesis. Furthermore, knockdown of CoRest in *Drosophila* males phenocopies the male infertility observed in LSD1 knockdown testes (Mačinković et al., 2019). The overlap in phenotypes between Lsd1 and CoRest mutants in the *Drosophila* spermatogenesis provides further evidence that *CoRest* may function with LSD1, but it is unclear if that function is in the germline, germline support cells, or both.

Analogous to the partial role for CoREST in *C. elegans* and mouse LSD1 maternal reprogramming, LSD1 may also be partially dependent on CoREST during mouse embryonic development. The phenotype of homozygous deletion of the *Lsd1* gene in mice is lethality by embryonic day 7.5 (e7.5) (Wang et al., 2007; Wang et al., 2009), while the *CoREST* deleted mice die by e16.5 (Yao et al., 2014). It is possible that the later embryonic lethality caused by loss of CoREST could result from the partial loss of LSD1 function, but this remains to be determined.

### Potential roles for CoREST in regulating LSD1 activity

There are two main possibilities for how CoREST may partially regulate LSD1 during maternal reprogramming in *C. elegans* and mice. One possibility is that CoREST is required for LSD1 activity at a subset of LSD1 targets. For example, it is possible that LSD1 needs CoREST to gain access to chromatin at certain targets that normally exist in more repressed chromatin. Consistent with this possibility, *in vitro* biochemical experiments showed that while LSD1 can demethylate H3K4 peptides or bulk histones, it is only capable of demethylating nucleosomes when in complex with CoREST (Lee et al., 2005; Shi et al., 2005; Yang et al., 2006). If CoREST is required for helping LSD1 gain access to certain chromatin targets, we would expect that the gene expression changes at these targets would also be completely affected by loss of CoREST. In contrast, at other genes where LSD1 does not need CoREST to gain access to chromatin, the loss of CoREST would not have the same effect as losing LSD1. However, we observe that most genes affected by the loss of LSD1 are also affected by the loss of CoREST, when sensitized by the loss of *met-2*. Furthermore, the gene expression changes caused by loss of CoREST are less affected than when LSD1 is lost. This is consistent with an alternative possibility, that CoREST helps LSD1 more efficiently access chromatin genome-wide. In this case, it is possible that SPR-5 maintains sufficient demethylase activity to prevent accumulation of H3K4me1/2 methylation in the absence of SPR-1. For this reason, mutation of SPR-1 alone would not result in a germline mortality phenotype. However, when MET-2 maternal reprogramming is lost, the inability to reprogram active chromatin states with H3K9me1/2 creates a chromatin environment where optimal SPR-5 activity is required. Without SPR-1, the reduced activity of SPR-5 is not sufficient to prevent the germline mortality phenotype that arises in *met-2; spr-1* double mutants. Thus, our data are more consistent with a model where CoREST is required maternally to help LSD1 more efficiently access chromatin genome-wide.

### Potential implications for CoREST function in humans

Taken together, our data in *C. elegans* and mice suggest that CoREST has a conserved role in maternal LSD1 reprogramming. The partial requirement for COREST in LSD1 function has potential implications for putative patients with mutations in *COREST*. The first human patients with *de novo* mutations in LSD1 have been identified. These patients display phenotypes that are similar to Kabuki Syndrome, which is characterized by developmental delay and craniofacial abnormalities (Chong et al., 2016; Tunovic et al., 2014). The *Lsd1* human mutations appear to be dominant partial loss of function mutations. It is possible that only partial loss of function mutants are viable because of the requirement for LSD1 in embryonic development and in stem cell populations (Haines et al., 2018; Kerenyi et al., 2013; Lambrot et al., 2015; Myrick et al., 2017; Tosic et al., 2018; Zhu et al., 2014). However, if CoREST is also required to help LSD1 more efficiently access chromatin genome-wide in humans, either maternally or zygotically, we might expect that loss of CoREST would readily give rise to similar developmental defects as those caused by partial loss of LSD1 function. As a result, we are actively searching for such potential human COREST patients.

## ACKNOWLEDGEMENTS

We are grateful to members of the Katz lab, as well as T. Caspary, T. Lee, and C. Bean, for their helpful discussion and critical reading of the manuscript; and the *Caenorhabditis* Genetics Center (funded by NIH P40 OD010440) for strains.

## Author Contributions

B.S.C., A.S. and D.J.K. conceived and designed the study and wrote the manuscript. B.S.C., A.S., R.G., S.R.C, M.C, and K.S. performed experiments under the direction of D.J.K. B.S.C, A.S., and D.J.K. analyzed data and interpreted results. D.A.M. helped with RNAseq analysis and visualizations. All authors discussed the results.

## Data availability

Raw and processed genomic data has been deposited with the Gene Expression Omnibus (www.ncbi.nlm.nih.gov/geo) under accession code GSE168081.

## MATERIALS AND METHODS

### *C. elegans* Strains

All *Caenorhabditis elegans strains* were grown and maintained at 20° C under standard conditions, as previously described (Brenner, 1974). The *C. elegans spr-5 (by101)(I)* strain was provided by R. Baumeister. The N2 Bristol wild-type (WT), *spr-1 (ar200)(V)*, and *et1*(III); *et1 [umnls 8 (myo-2p :: GFP + NeoR, III: 9421936)](V)* strain was provided by the *Caenorhabditis* Genetics Center. The *met-2 (n4256)(III) strain* was provided by R. Horvitz. From these strains we generated *spr-1 (ar200) (V)*/ *et1 [umnls 8 (myo-2p :: GFP + NeoR, III: 9421936)*]*(V)* and *met-2 (n4256) (III)* / *et1 [umnls 8 (myo-2p :: GFP + NeoR, III: 9421936)](V); spr-1 (ar200)(V)*/ *et1 [umnls 8 (myo-2p :: GFP + NeoR, III: 9421936)](V)*. For genotyping, single animals were picked into 5-10ul of lysis buffer (50mM KCl, 10mM Tris-HCl (pH 8.3), 2.5mM MgCl_2_, 0.45% NP-40, 0.45% Tween-20, 0.01% gelatin) and incubated at 65°C for 1 hour followed by 95°C for 30 minutes. PCR reactions were performed with AmpliTaq Gold (Invitrogen) according to the manufacturer’s protocol and reactions were resolved on agarose gels.

### Generation of M448V hypomorphic allele

Oligos were designed to include an A>G SNP conversion which removed an HpyAV restriction site, and a G>A PAM blocking silent SNP. C57BL/6 females were superovulated by injecting 0.1mL/head of CARD HyperOva (i.p.) on day 1. After 48 hours, females were injected with 7.5IU human chorionic gonadotropin (hCG, i.p.). Oocytes were collected 13 hours after the administration of hCG and fertilized with C57BL/6 sperm in vitro. Five hours postfertilization, 50ng/uL Cas9mRNA, 50ng/uL oligo, and 50ng/uL sgRNA were injected into the cytoplasm of embryos. Injected embryos were incubated at 37°C overnight. Two-cell embryos were then transferred into the oviducts of pseudopregnant females. Progeny of those females were genotyped for the point mutation, mated, and progeny were genotyped again to ensure the mutation passed through the germline. Mutant animals were backcrossed at least s two times to C57BL/6 animals before being used in experiments.

### Mouse husbandry and genotyping

The following mouse strains were used: *Zp3-Cre* MGI:2176187 (de Vries et al. 2000), *Lsd1*^*fl/fl*^ MGI: 3711205 (Wang et al. 2007), C57BL/6 MGI: 3715241, and *Lsd1*^*M448V*^. Primers for Lsd1 forward (F): GCACCAACACTAAAGAGTATCC, Lsd1 reverse (R): CCACAGAACTTCAAATTACTAAT. A wild type allele of Lsd1 results in a 720 base pair (bp) product, the floxed allele is 480 bp, and the deleted allele is 280 bp. Primers for Cre F: GAACCTGATGGACATGTTCAGG, Cre R:

AGTGCGTTCGAACGCTAGAGCCTGT, Cre ctrl F: TTACGTCCATCGTGG ACAGC Cre ctrl R: TGGGCTGGGTGTTAGCCTTA. If Cre+, this results in a 302 bp product, and Cre ctrl F/R primers are an internal control that yields a 250bp product. Primers for M448V F: CCCAAATGGCATGACATAAA, M448V R: TAAGGCACCAAACCCCTTCT resulting in a 386bp product. The point mutation removes a restriction site, so mutants versus wild type were determined by incubating PCR products at 37°C with the HpyAV restriction enzyme for one hour. Wild type band sizes: 72bp, 81bp, 209bp, 24bp. M448V band sizes: 72bp, 290bp, 24bp. All mouse work was performed under protocols approved by the Emory University Institutional Animal Care and Use Committee.

### Immunofluorescence

Mice were sacrificed by cervical dislocation and ovaries were isolated. Ovaries were then fixed in 4% PFA for one hour, followed by four PBS washes over two hours. Tissues were cryoprotected in 30% sucrose at 4°C overnight and then embedded in O.C.T. Compound (Tissue Tek). Cryosections were obtained at 10µm and immunostaining was performed using rabbit polyclonal anti-LSD1 (1:200, ab17721), rabbit polyclonal anti-CoREST (1:100, LS-B8140-50), and Alexa fluor conjugated secondary antibodies (1:500).

### Perinatal lethality

Breeding cages were observed daily for new litters and number born alive were scored at P0. At P1, litter sizes were scored again, and percent lethality was calculated by determining the number of animals that died divided by the original size of the litter. Those that died due to failure to thrive shortly after birth were often missing visible milk spots. Only litters from mothers <8 months of age were used to avoid complications due to advanced maternal age.

### Germline mortality assay

The germline mortality experiments were performed as described by Katz and colleagues (Katz et al., 2009). In brief, worms were maintained at 20° C and three fertile young adults with visible embryos were transferred to new NGM plates every four days. The total number of progeny from wild type, *spr-1* mutants, and *spr-5* mutants was counted every third generation until generation 17, after which counts were completed every other generation. Average number of progeny from *spr-5* mutants was calculated from 10 animals until counts were stopped at generation 41 due to the inability to maintain fertile animals. For wild type, the average number of progeny was calculated from five animals until generation 41 when the average number of progeny was calculated from six animals. The average number of progeny from *spr-1* mutants was calculated from ten animals throughout the entirety of the experiment. The same germline morality assay was adapted to evaluate the germline mortality of wild-type, *spr-1* and *met-2* single mutants, and *met-2; spr-1* double mutants. Here, the number of progeny were counted every generation, except from *spr-1* mutants which was counted every fourth generation. The average number of progeny from wild type, *spr-1* mutants and *met-2* mutants was calculated from ten animals, while *met-2; spr-1* was calculated from 30 animals. The standard error of the mean (SEM) was calculated for each generation the number of progeny were averaged.

### RNA sequencing and analysis

Total RNA was isolated using TRIzol reagent (Invitrogen) from ~500-1000 starved L1 larvae hatched at room temperature (21°C −22°C) overnight in M9 Buffer. L1 larvae from wild type, *spr-1, met-2*, and *met-2; spr-1* were isolated at generation 7 (F7) prior to the observed decrease in sterility. For each genotype, 2 biological replicates were obtained. Sequencing reads were checked for quality using FastQC (Wingett and Andrews, 2018), filtered using Trimmomatic (Bolger et al., 2014), and remapped to the *C. elegans* transcriptome (ce10, WS220) using HISAT2 (Kim et al., 2015a). Read count by gene was obtained by FeatureCounts (Liao et al., 2014). Differentially expressed transcripts (significance threshold, Wald test, p-value < 0.05) were determined using DESEQ2 (v.2.11.40.2) (Love et al., 2014). Transcripts per million (TPM) values were calculated from raw data obtained from FeatureCounts output. Subsequent downstream analysis was performed using R with normalized counts and p-values from DESEQ2 (v.2.11.40.2). Heatmaps were produced using the ComplexHeatmap R Package (Gu et al., 2016). Data was scaled and hierarchical clustering was performed using the complete linkage algorithm. In the linkage algorithm, distance was measured by calculating pairwise distance. Volcano plots were produced using the EnhancedVolcano package (v.0.99.16). Additionally, Gene Ontology (GO) Pathway analysis was performed using the online platform WormEnrichr (Chen et al., 2013; Kuleshov et al., 2016). An additional heatmap comparison of differentially expressed genes between *spr-1, met-2, met-2; spr-1* and *spr-5; met-2* progeny compared to wild type progeny was generated in Microsoft Excel using log2 fold change values from the DESEQ2 analysis. Differentially expressed genes in *spr-5; met-2* double mutants compared to wild type examined in this manuscript were obtained from a separate RNAseq analysis performed under the same conditions (Carpenter et al., 2021). Because transcript isoforms were ignored, we discuss the data in terms of “genes expressed” rather than “transcripts expressed”.

### Differential interference contrast (DIC) microscopy

Worms were immobilized in 0.1% levamisole and placed on a 2% agarose pad for imaging at either 10x, 40x, or 100x magnification.

## Supplemental Material

**Figure S1.**
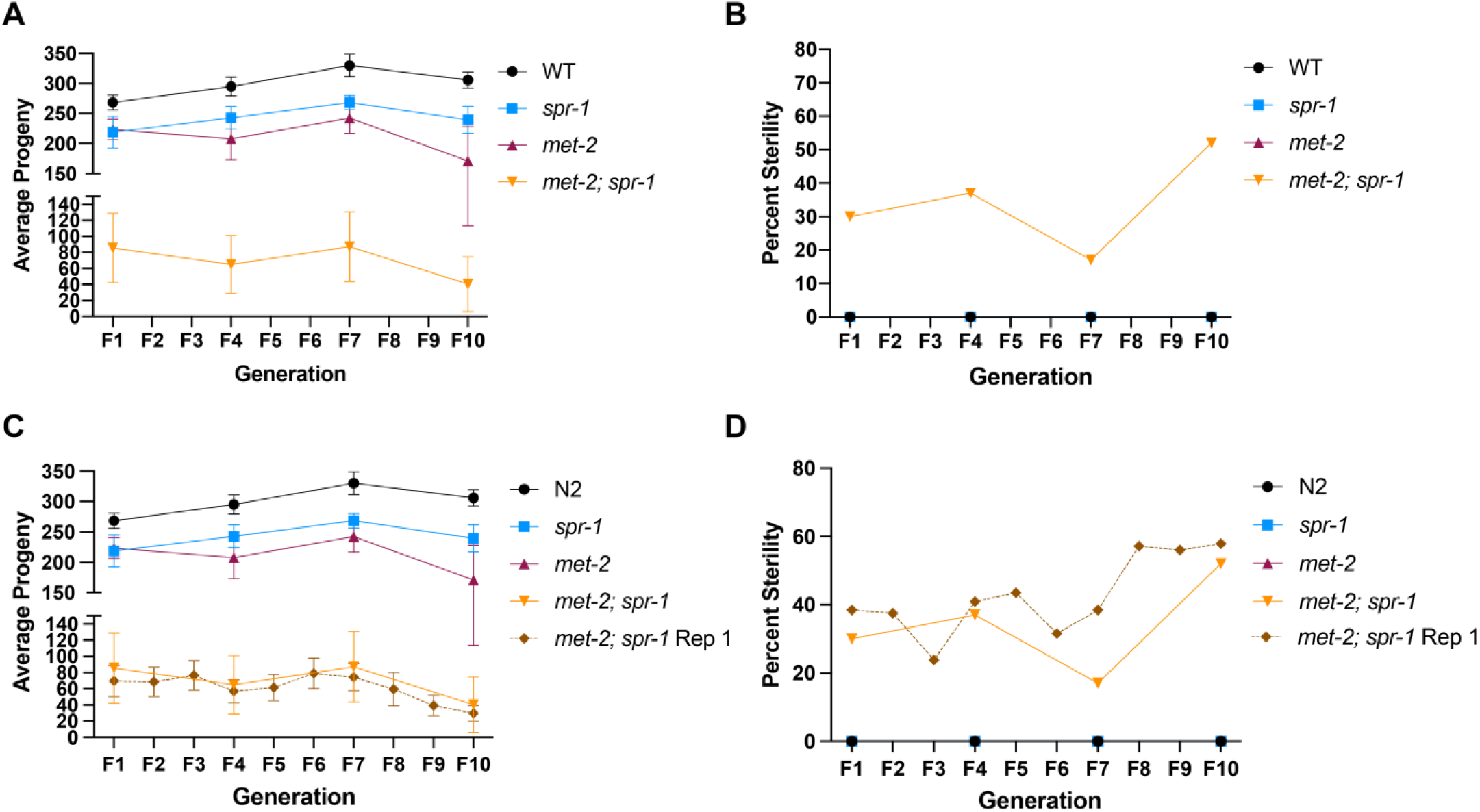
Germline mortality in *spr-1* and *met-2; spr-1* mutants replicate experiment. (A) The average number of total progeny from wild type (WT), *spr-1, met-2* and *met-2; spr-1* mutants over progressive generations in a repeat experiment (see Fig. 1E for comparison). (B) Percent of animals cloned out for repeat experiment in (A) scored for sterility over progressive generations (see Fig. 1F for comparison). (C) Same as (A) with overlay of total progeny from *met-2; spr-1* (*met-2; spr-1* Rep 1) mutants from germline mortality experiment in Figure 1E. (D) Same as (B) with overlay of percent sterile *met-2; spr-1*(*met-2; spr-1* Rep 1) mutants in Figure 1F. Error bars in (A, C) represent the standard error of the mean (SEM). Animals cloned out for the replicate germline mortality experiment were only scored at F1, F4, F7, and F10 generations in (A-D).

**Figure S2.**
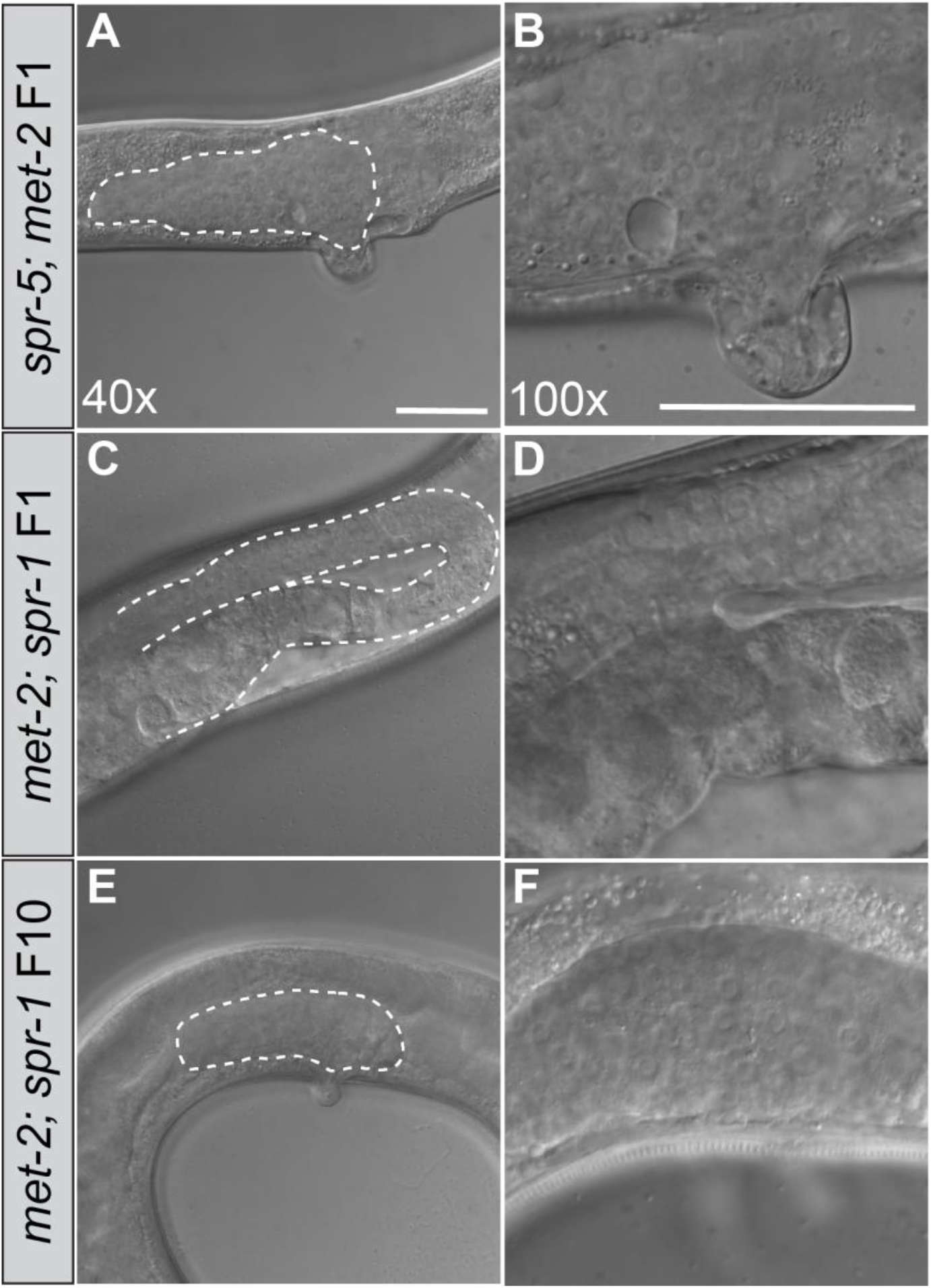
Sterile *spr-5; met-2* and *met-2; spr-1* double mutant gonads. 40x Differential Interference Contrast (DIC) images of F1 *spr-5; met-2* (A) *F1 met-2; spr-1* (C), and F10 *met-2; spr-1* (E) adult germlines. White dotted lines denote germlines. 100x DIC images of F1 *spr-5; met-2* (B) F1 *met-2; spr-1* (D), and F10 *met-2; spr-1* (F) adult germlines. Scale bar: 100μm.

**Figure S3:**
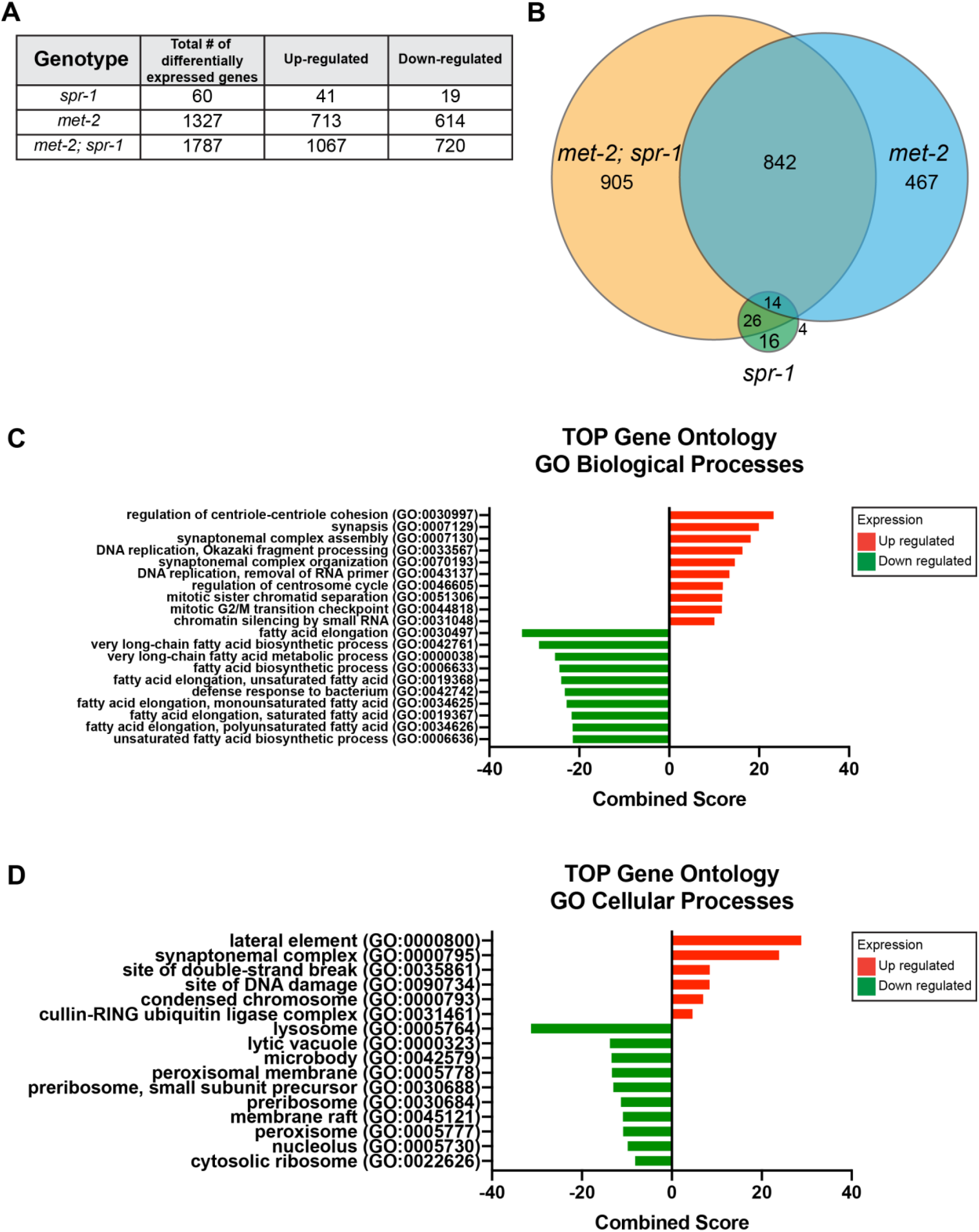
Differential gene expression *spr-1, met-2*, and *met-2; spr-1* progeny compared to wild type. (A) Table summary of differentially expressed genes (DEGs) in *spr-1, met-2, and met-2; spr-1* progeny from DeSEQ2 analysis (significance cut-off of p-adj< 0.05). (B) Overlap of differentially expressed genes between *spr-1, met-2, and met-2; spr-1* L1 progeny. Gene Ontology analysis showing Biological Processes (C) and Cellular Components (D) amongst genes that were up-regulated (red) and down-regulated (green) in *met-2; spr-1* progeny compared to wild-type. Combined Score was computed to determine gene category enrichment (Chen et al., 2013).

**Figure S4:**
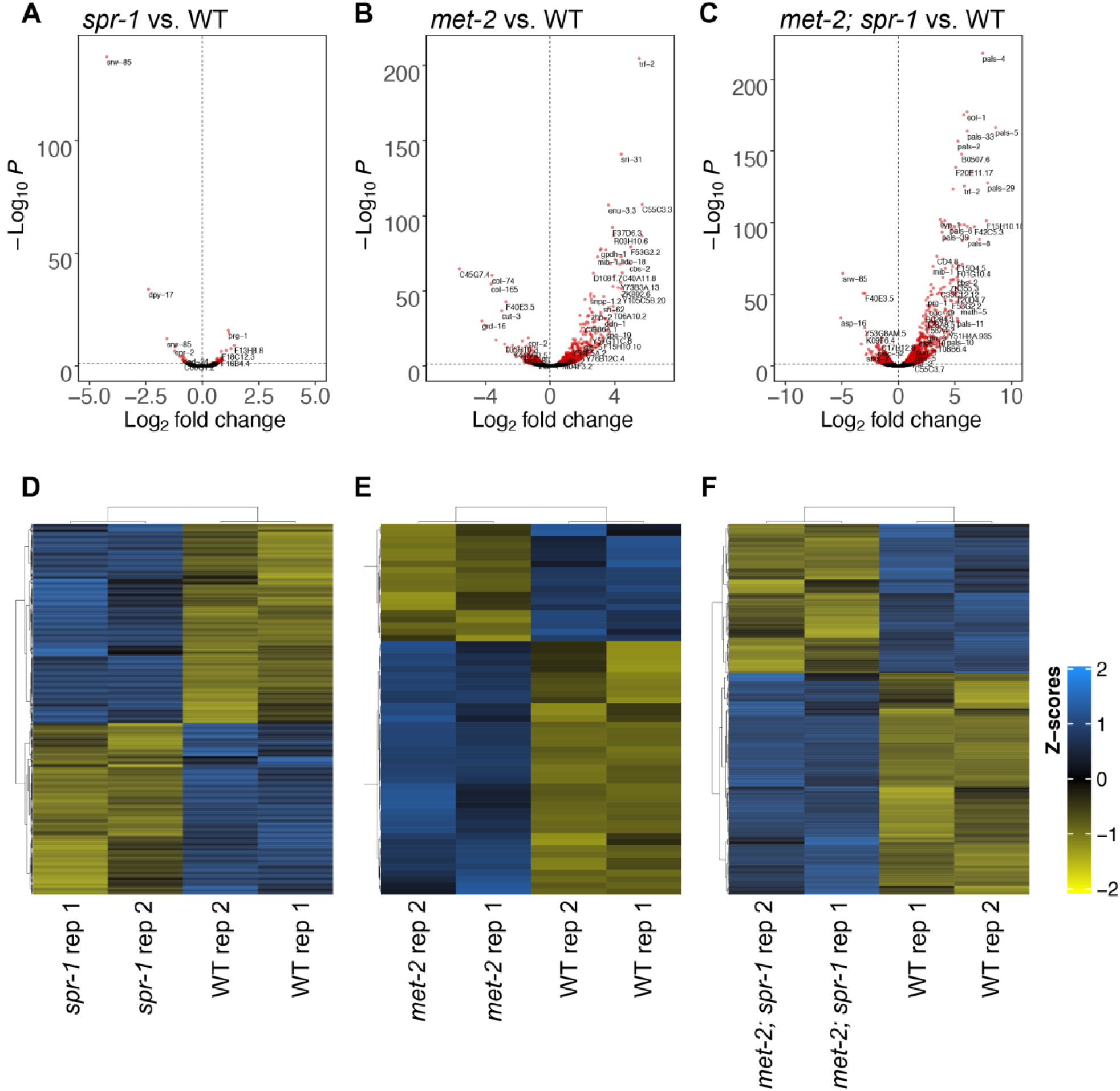
Differential expression and replicate comparison of RNAseq experiments performed on wild type, *spr-1, met-2*, and *met-2; spr-1* progeny. Volcano plot of log2 fold changes in gene expression (x-axis) by statistical significance (-Log_10_ P-value; y-axis) in *spr-1* (A), *met-2* (B), and *met-2; spr-1* (C) L1 Progeny compared to wild type (WT). Heatmap of differentially expressed RNA-seq transcripts between WT and *spr-1* (D), *met-2* (E), and *met-2; spr-1* (F). Data was scaled and hierarchical clustering was performed using complete linkage algorithm, with distance measured by calculating pairwise distance. Higher (blue) and lower (yellow) expression is reported as a z-score.

**Figure S5.**
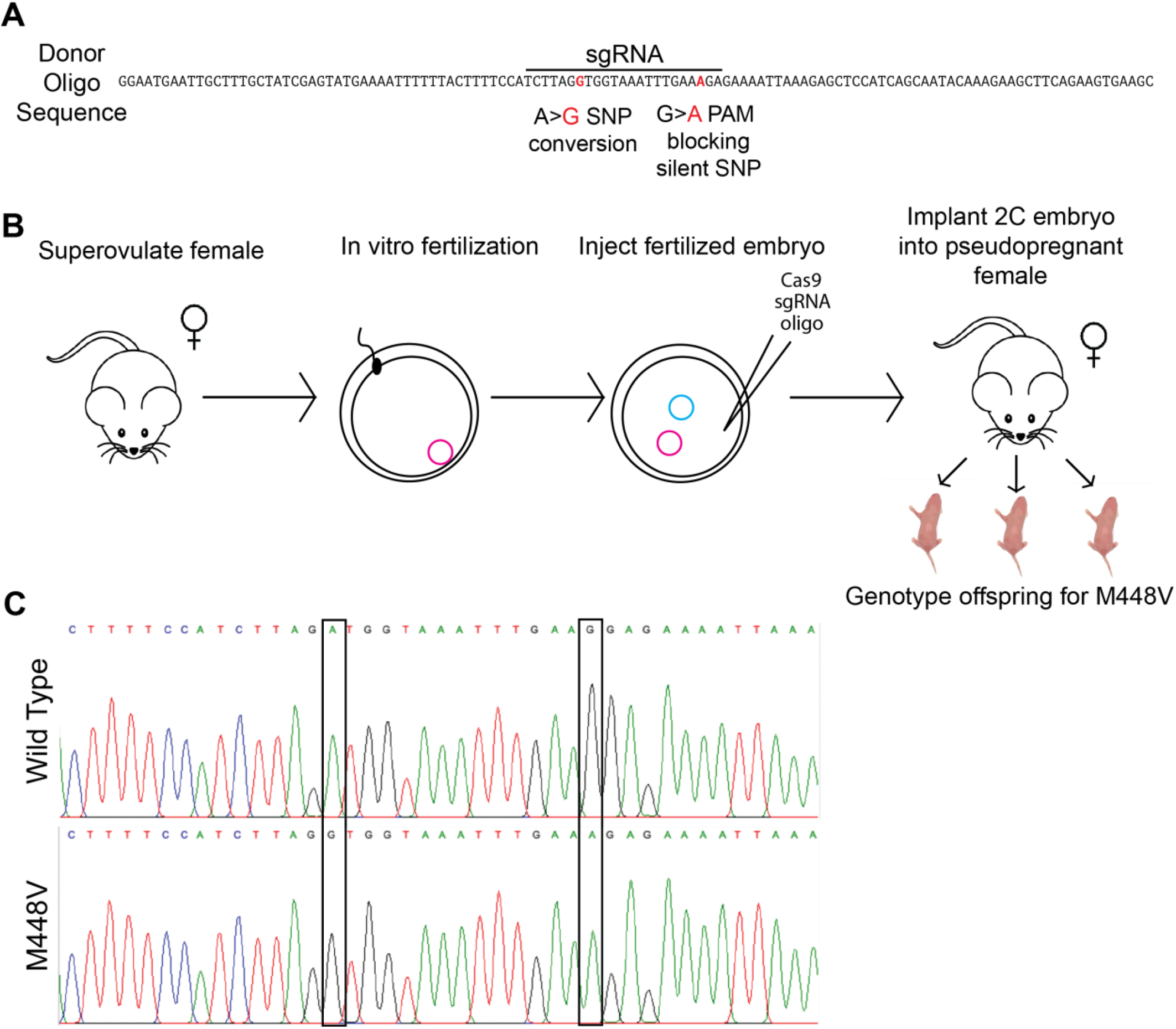
Generation of M448V hypomorphic allele. (A) Donor oligo sequence with the sgRNA sequence denoted. Two point mutations being introduced are colored in red: A>G SNP and G>A PAM blocking silent SNP. (B) Workflow for introducing M448V mutation into mice. (C) Wild-type chromatogram versus validated A>G SNP and G>A silent SNP in the newly generated mutant.

**Figure S6.**
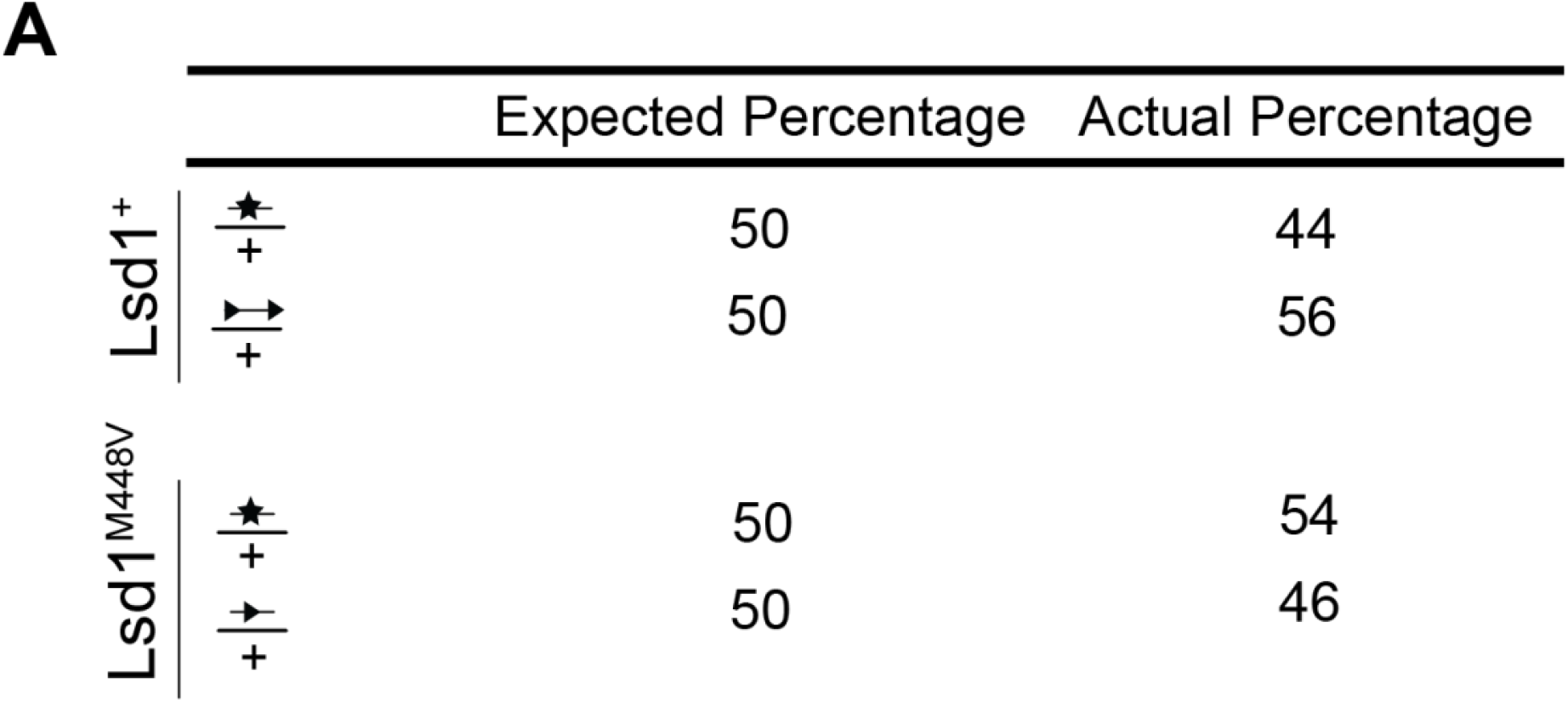
Percent survival by genotype per experimental condition. *Lsd1*^*+*^ n=50 animals genotyped. *Lsd1*^*M448V*^ n=87. Statistical analyses performed using chi square test, no significant difference between expected and actual percentages for either experimental condition.

## Supplemental files

**Supplemental file 1**. List of individual litters analyzed for embryonic lethality in Figure 5.

